# Imaging of Morphological and Biochemical Hallmarks of Apoptosis with Optimized Optogenetic Actuators

**DOI:** 10.1101/551788

**Authors:** Walton C. Godwin, George F. Hoffmann, Taylor J. Gray, Robert M. Hughes

## Abstract

The creation of pathway-specific optogenetic switches for the activation of cell death pathways can provide insight into the mechanisms of apoptosis while forming a basis for non-invasive, next generation therapeutic strategies. Previous work has demonstrated the utility of Cryptochrome 2 (Cry2)/CIB, a blue light activated protein – protein dimerization module from A. thaliana in conjunction with BAX, an OMM targeting pro-apoptotic protein, for light-mediated initiation of mitochondrial outer membrane permeabilization (MOMP) and downstream apoptosis. In this work, we have further developed the original light activated Cry2-BAX system (henceforth referred to as “OptoBAX”) by improving the photophysical properties and light-independent interaction of our optogenetic switch. The resulting optogenetic constructs have significantly reduced the frequency of light exposure required for the activation of membrane permeabilization, in addition to reducing dark state cytotoxicity. Furthermore, we utilize OptoBAX in a series of experiments designed to measure the timing of the dramatic morphological and biochemical changes that occur in apoptotic cells following light-induced permeabilization of the outer mitochondrial membrane. Utilizing this data, we construct a timeline of biochemical and morphological events in early apoptosis, in addition to demonstrating a direct link between MOMP-induced redistribution of actin and apoptotic progression.

**Significance Statement:** Apoptosis, the cycle of programmed cell death, is a critical process in organism development. Unraveling the complex biochemical events that drive apoptotic progression remains an ambitious goal in cell signaling research. To this end, we describe the development and optimization of optogenetic proteins that initiate the apoptotic pathway in mammalian cells by recruiting BAX, a 21-kD pro-apoptotic Bcl-2 family protein, to the outer mitochondrial membrane with light. We utilize these optogenetic constructs in tandem with fluorescent reporter molecules for imaging key events in early apoptosis, including membrane inversion and caspase cleavage, in addition to tracking the redistribution of actin, a key protein for apoptotic progression. This work provides insight into the relative timing and interplay between early events in apoptosis.

## INTRODUCTION

Among the wide-ranging applications of optogenetic proteins – which include light-mediated control of transcription[1], ion channels[2], and enzymatic activity[3] – the use of light for wholesale activation or inhibition of cell signaling pathways presents a uniquely powerful approach for the determination of cell fate – acting as a rapid, non-invasive trigger for cascades of biochemical events that ultimately decide whether a cell proliferates or dies. After many years of effort, numerous cell signaling pathways critical for cell fate decisions have been elucidated, including pathways for the control of cell division[4], cell motility[5], and apoptosis[6]. As a result, it is possible to identify individual proteins that serve as highly specific effector biomolecules for the initiation of signaling cascades undergirding these vital processes. Such effector molecules are ideal candidates for incorporation into light-activated protein switches for control of cell signaling pathways[7–9]. These switches are useful for the study of not only the effector molecules themselves, but also of biochemical events that occur in the wake of their activation. In the case of apoptosis, which can be viewed as a singular destination (“cell death”) attainable by various routes[10], numerous effector molecules exist that could serve as candidates for optical control of apoptosis, including the Bcl-2 family proteins[11]. Bax, a 21-kD protein which is a key effector of mitochondrial membrane permeabilization (MOMP) in the intrinsic apoptosis pathway, is one such effector molecule with an in vivo activity profile that makes it ideal for incorporation into an optogenetic switch: phosphorylation-gated activity, predominantly cytosolic localization in the “off” state, and robust initiation of permeabilization of the outer mitochondrial membrane (OMM)[12–15]. In previous work, we have demonstrated that the incorporation of BAX into a cryptochrome 2 (Cry2) – cryptochrome interacting β helix-loop-helix (CIB) based optogenetic switch enables light activated MOMP and release of apopotic effector proteins such as Smac-1 [16]. A key step in the engineering of this switch was the implementation of a phospho-mimetic mutation in the Bax C-terminus at serine 184 (S184E), which maintains Bax in a predominantly cytosolic state by increasing the off-rate of localization at the OMM, thus eliminating one potential source of light-independent background[17]. However, another potential source of light-independent (or “dark”) background for this optogenetic switch, is the dark state interaction of the Cry2 photoreceptor with the mitochondrial Tom20-CIB construct. In this work, we have created next generation iterations of the Cry2-CIB Bax system (“OptoBax”), which we refer to as “OptoBax 2.0”. The optimized versions of this cell death switch include longer Cry2 variants designed to reduce the dark state interaction with CIB, and incorporate a mutation that extends the photocycle of the Cry2-CIB interaction, promoting longer association times with the OMM[18]. This optimized construct requires less frequent light stimulation to maintain OptoBAX and the OMM, and advantage for long timecourse experimentation where blue light toxicity may be of concern[19]. In addition, while the initial studies with OptoBax were conducted in HeLa cells, we have ported the OptoBax 2.0 system into the cell lines Neuro-2a and HEK-293T, demonstrating light initiated apoptosis in these cell lines analogous to that previously observed in HeLa cells. Finally, in HEK-293T cells, we use a combination of fluorescently-tagged proteins and commercially available dyes for multi-color monitoring of critical steps in apoptosis, downstream of initial mitochondrial insult (cytoskeletal rearrangement, chromatin condensation, membrane inversion, and caspase cleavage), and compare the our results to those previously observed with a classic small molecule initiator of the apoptotic cascade. We use the resulting data to create a timeline of key apoptotic events in HEK-293T cells, in addition to providing insight into changes in actin localization and degradation of the nuclear/cytoplasmic barrier, both key steps in early apoptosis.

## Results and Discussion

### Creation and validation of new optogenetic constructs

Initial versions of the OptoBax system were comprised of BAX as a fusion to the N-terminus of a Cry2PHR (1-498)-mCherry (BAX.Cry2PHR.mCh) fusion protein or as a fusion to the C-terminus of a Cry2PHR (1-498)-mCherry fusion (Cry2PHR.mCh.BAX), in conjunction with the Cry2-interacting partner CIB in fusion with a Tom20 OMM localization domain with or without a C-terminal GFP (Tom20.CIB.GFP or Tom20.CIB)[20]. While both BAX fusions were demonstrated to initiate light-mediated apoptosis in HeLa cells, the C-terminal fusion of a BAX mutant (S184E) was demonstrated to be an effective light-activated apoptotic switch, with reduced association of BAX with the OMM due to the phospho-mimic S184E, and a more rapid induction of the apoptotic cascade than the N-terminal BAX construct[16]. Subsequent to the completion of these initial studies, optimized versions of the Cry2 optogenetic system were reported, demonstrating, in the context of a yeast two-hybrid assay, that extending the truncated Cry2PHR from 498 to 515 amino acids or longer in length (up to 535 aa) significantly reduced the light independent background present in the optogenetic system [18]. In the same report, a genetic screen also identified a Cry2 mutant (L348F) that extends the photocycle of Cry2 well beyond that of the wild type protein, prolonging the half-life of the light-promoted Cry2-CIB interaction from ~6 min to ~24 min[18]. This mutation prolongs the lifetime of the light-promoted semiquinone form of FAD, the light responsive Cry2 cofactor. We hypothesized that incorporating similar changes into OptoBax might result in an improved pro-apoptotic switch, via reduced light-independent Cry2/CIB interaction, and, with incorporation of the L348F mutant, reduced light input than the original OptoBax, creating a system more easily amenable to incorporation into drug discovery platforms or into model organisms. As a result, for the OptoBax 2.0 system, we created two additional Cry2–Bax fusions: Cry2(1-531).mCh.Bax.S184E and Cry2(1-531).L348F.mCh.Bax.S184E, pairing them with the original Tom20.CIB.GFP and Tom20.CIB constructs (Fig. 1A). We subsequently performed tests of light-independent cell death (“dark background”) in dually-transfected HeLa cells with a fluorescent indicator of cell viability (Supporting Fig. 1). We found that, versus the original OptoBax system, dark background was reduced from ~36% (Cry2PHR.mCh.Bax.S184E) to ~20% (Cry2(1-531).mCh.Bax.S184E and Cry2(1-531).L348F.mCh.Bax.S184E), similar to background cell death observed with the Cry2.mCh fusions without Bax. We further validated the impact of the L348F mutant in the Cry2.mCh fusions in HeLa cells, utilizing a single pulse of 470 nm light to promote association of Cry2 with the OMM-localized CIB domain (Fig. 1B-C and Supporting Movie 1), confirming that the L348F mutant extended the half-life of association with the OMM from 6.25 (+/-0.5) min to 32.8 (+/-1.0) min. We also investigated whether the Cry2(1-531).L348F.mCh.BAX.S184E construct was also an efficient inducer of apoptosis, versus the Cry2(1-531).L348F.mCh fusion (Fig. 2A and Supporting Movie 2), in addition to investigating whether the L348F mutant was able to induce apoptosis with less frequent light stimulation (Fig. 2B). We designed an experiment which set the blue light illumination interval at 10 min, which is longer than the half-life of the WT construct and significantly shorter than that of the extended photocycle mutant. As a consequence of this experimental design, the non-mutant Cry2 construct will repeatedly revert to the cytosolic state during the course of the experiment, while the L348F mutant will maintain consistent localization at the OMM. We counted the number of apoptotic cells (based on the loss of adherent cell morphology) after two hours of simultaneous blue light stimulation and imaging on a Leica Dmi8 widefield fluorescent microscope. In this study, 68% of transfected cells (28/41 total cells) expressing the Cry2(1-531).L348F.mCh.BAX.S184E/Tom20.Cib.GFP fusion became apoptotic vs. 20% of transfected cells (11/55 total cells) expressing the standard photocycle Cry2(1-531).mCh.BAX.S184E construct, indicating that prolonging the photocycle enables more apoptosis with less frequent light input. We also observed some background apoptosis with the long-photocycle fusion without Bax (Cry2(1-531).L348F.mCh; 3/59 total cells). This is likely due to loss of mitochondrial polarization due to sustained recruitment of the long-photocycle mutant, as no background apoptosis was observed with its standard half-life analogue (Cry2(1-531).mCh; 0/48 total cells).

**Figure 1.**
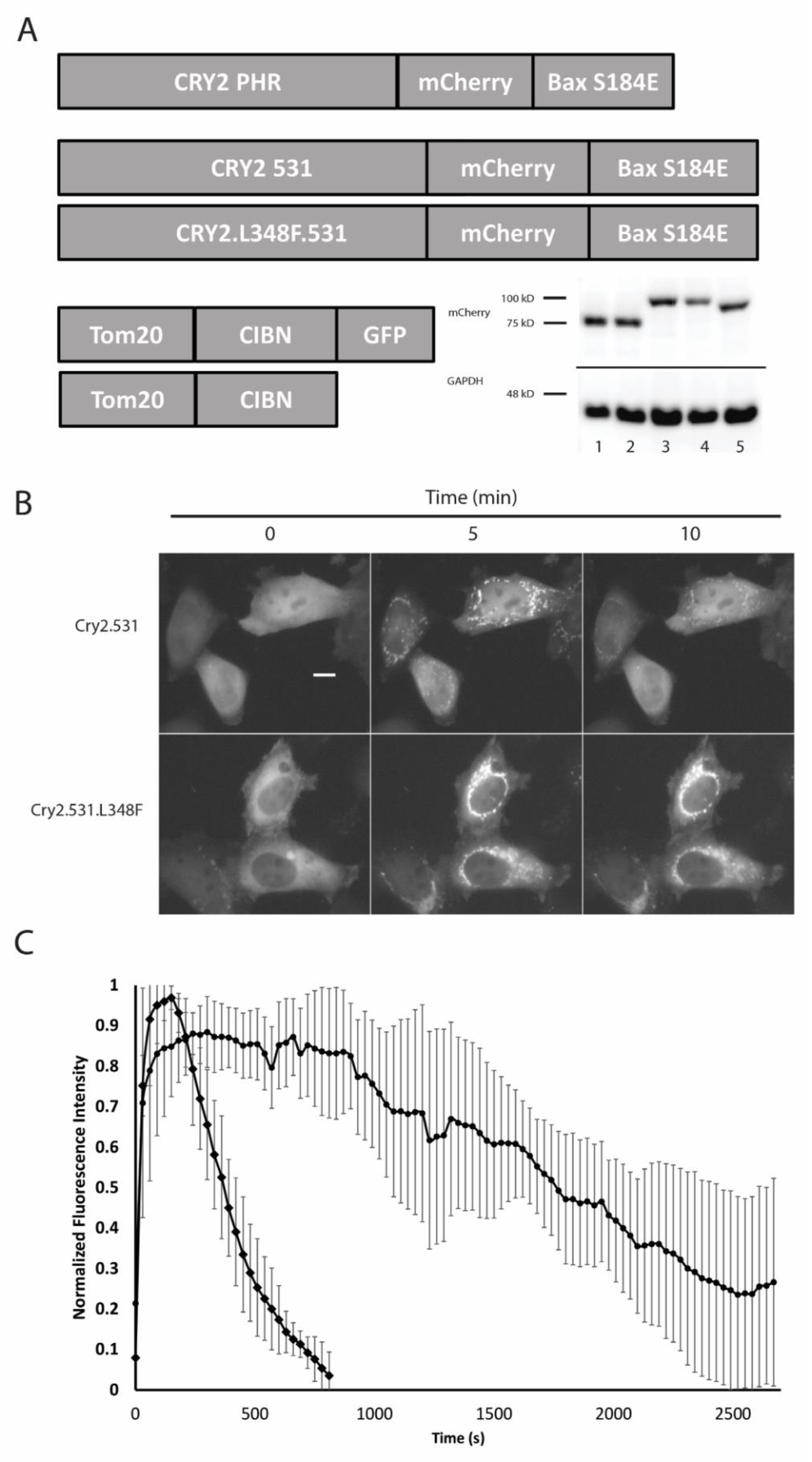
REDESIGN OF OPTOGENETIC ACTUATORS AND CHARACTERIZATION OF EXTENDED PHOTOCYCLE MUTANT L348F. A. Optimized versions of OptoBax created for reduction of light-independent cell death and extension of photocycle. Western blot (α-mCherry; inset) lanes are (1) Cry2.531.mCherry; (2) Cry2.531.L348F.mCherry; (3) Cry2.531.mCherry.Bax; (4) Cry2.531.L348F.mCherry.Bax; (5) Cry2PHR.mCherry.Bax. B. Demonstration of elongated photocycle in the Cry2 L348F mutant. Images shown are before, 5 min, and 10 min post activation with a 150 ms pulse of 470 nm light. C. Plot of normalized fluorescence data from the experiment shown in 1.B. Scale bar = 10 microns.

**Figure 2.**
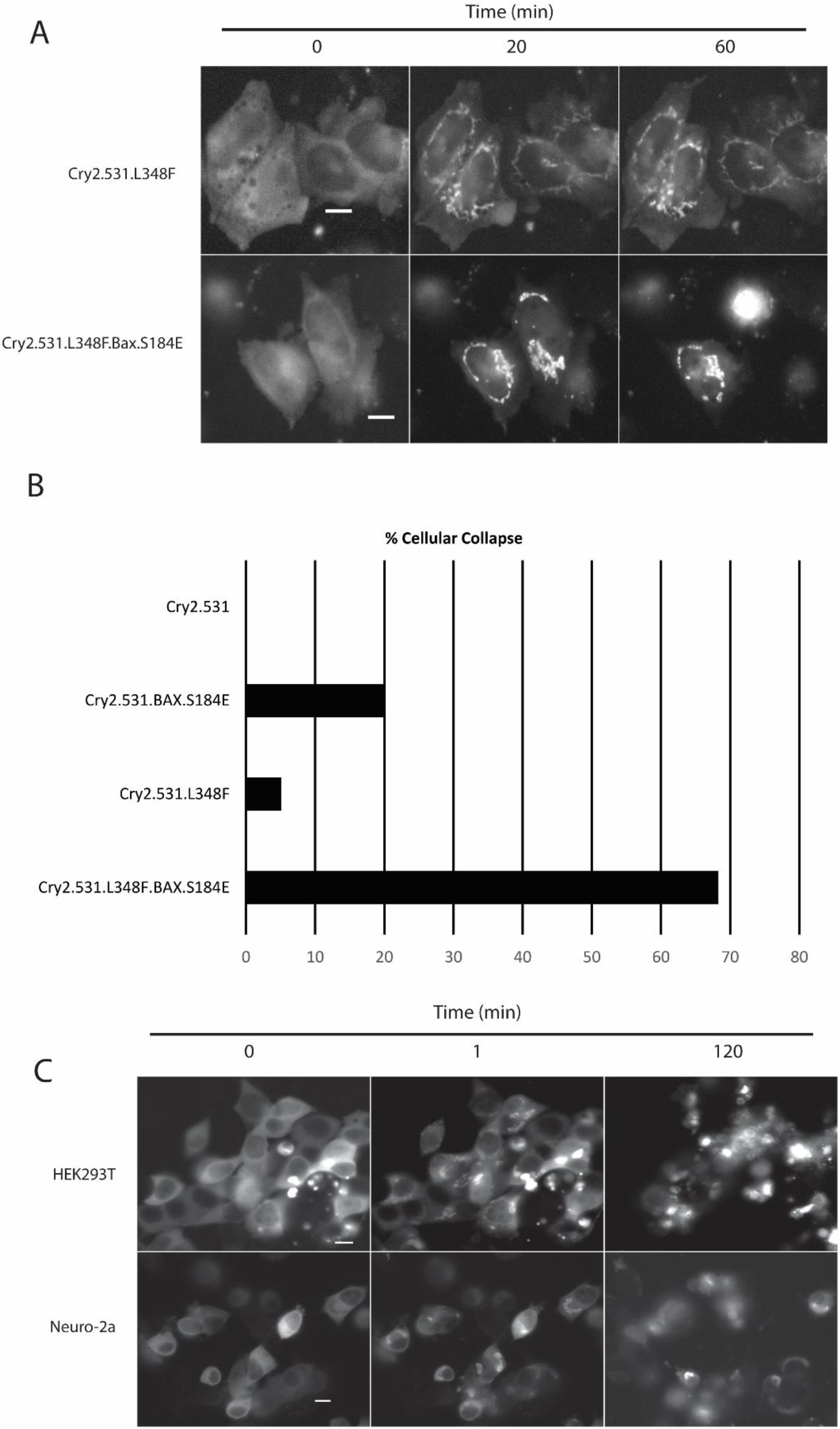
MINIMAL LIGHT APPLICATION REQUIRED FOR APOPTOTIC CELL DEATH AND APPLICATION TO OTHER CELL LINES. A. OMM recruitment of OptoBax constructs with (Cry2.531.L348F.Bax/ Tom20.Cib.GFP) and without (Cry2.531.L348F/ Tom20.Cib.GFP) the proapoptotic Bax fusion in HeLa cells. Cells illuminated with a 150 ms pulse of 470 nm light every 10 minutes over a two hour time course. B. Quantification of cellular collapse over the two hour time course experiment. C. Demonstration light induced cellular collapse with OptoBax (Cry2.531.L348F.Bax/Tom20.Cib.GFP) in HEK293T and Neuro-2a cell lines over a two hour time course (470 nm; 150 ms pulse; images acquired every minute). Scale bar = 10 microns.

### Translation to other cell lines

Given the common mechanism of Bax-mediated MOMP and subsequent apoptosis in mammalian cells[21], we anticipated that our optogenetic system might readily translate to other mammalian cell lines. In order to test this assumption, we deployed the OptoBax 2.0 constructs in two additional cell lines, Neuro-2a and HEK293T (Fig. 2C). Similar results were obtained in both cell lines, with induction of apoptosis upon recruitment of the Cry2(1-531).L348F.mCh.BAX.S184E to the OMM with light. In particular, the reliably high transfection efficiency observed with the HEK293T cell line made further exploration of the apoptotic cascade in this cell line particularly appealing. As a result, we carried out the remainder of our experiments in the HEK293T cell line.

### Tracking the hallmarks of apoptosis with optogenetic techniques

The morphological and biochemical hallmarks of apoptosis include cell blebbing, loss of adhesion and cellular volume, chromatin condensation and nuclear fragmentation, cytoskeletal rearrangement, phosphatidylserine inversion, release of mitochondrial proteins (Cytochrome C/Smac/Diablo), and caspase cleavage[22]. Fluorescently labelled proteins and small molecules for tracking these key apoptotic events have become widely available in recent years[23–26]. This makes practical the prospect of using a light activated system for the induction of apoptosis in conjunction with these live cell compatible markers on a multi-channel fluorescent microscope equipped with a live cell incubator. One potentially useful application of the OptoBax system, in conjunction with the forementioned markers, is the establishment of a timeline of key apoptotic events, in order to define not only the order, but also the interplay between key steps in the apoptotic cascade. In turn, the establishment of a timeline around a specific event (i.e., Bax translocation to the OMM) may provide novel insights into their interdependence. To this end, we selected several markers, including a nuclear stain (Hoescht 33342) for monitoring chromatin condensation, a marker of phosphatidylserine (PS) inversion (Annexin V-FITC & Annexin V-Cy5), a cytoskeletal marker (GFP-Actin [27]), and a marker for downstream caspase cleavage (CellEvent Caspase 3/7). In our first experiment, we sought to simultaneously monitor chromatin condensation, cytoskeletal rearrangement, and phosphatidylserine inversion in the wake of BAX translocation to the OMM. In the HEK293T cell line, we were able to efficiently transfect both components of the OptoBax 2.0 system (Cry2(1-531).L348F.mCh.Bax.S184E and Tom20.CIB), in addition to a third protein, a cytoskeletal marker for actin (GFP-Actin). In the same experiment, we labeled chromatin with Hoescht 33342 in addition incorporating an Annexin V-Cy5 fusion as a marker for PS inversion. This provided a 4-color system for monitoring light induced BAX translocation to the mitochondria (mCherry/TxRed), cytoskeletal rearrangement (FITC/GFP), chromatin condensation (DAPI), and membrane flipping (Cy5). We subsequently measured the impact of BAX-mediated apoptosis on the cytoskeleton and chromatin over the course of a 2 hour illumination experiment (Fig. 3 and Supporting Movie 3). As anticipated, we observed changes in the localization of our GFP-Actin construct, indicative of cytoskeletal rearrangement, nuclear collapse and intensification of the fluorescence intensity of the Hoescht 33342 dye, indicative of chromatin condensation, and incorporation of the Annexin V-Cy5 dye, indicative of phosphatidylserine inversion (Fig. 3A). We were also able to observe redistribution of the GFP-Actin marker from the cytosol to both the nucleus and the mitochondria (44% (111/252) cells analyzed clearly demonstrated mitochondrial GFP-Actin 70 min post-OptoBax activation; Fig. 3B-3C). The accumulation of actin at the mitochondria remains a poorly understood event in the BAX-mediated apoptosis pathway[28–31], while the accumulation of actin and numerous actin-binding proteins (cofilin 1; CAP1) at the OMM in the wake of small molecule induced apoptosis is often characterized as an early event that occurs well prior to BAX translocation[29, 30, 32]. In our system, however, membrane permeabilization with OptoBAX precipitates downstream actin redistribution to the OMM well after BAX recruitment (~30 min post-OptoBAX activation). This surprising result points to a closer relationship between cytoskeletal dynamics, Bcl-2 family translocation events, and apoptotic progression than previously described[28]. As a follow up to this initial result, we repeated our multi-color experiments in the presence of compounds known to have opposing effects on actin dynamics: jasplakinolide, which inhibits actin depolymerization, maintaining the F-actin state, and latrunculin A, which inhibits actin polymerization by sequestering G-actin[33]. As expected, treatment of cells with these compounds resulted in strikingly different distributions of our actin-GFP reporter (Supporting Figure 2). In addition, both compounds delayed the entry of cells into light-activated apoptosis in comparison to the no treatment control, as assessed by both the reduction of chromatin condensation and PS externalization during a two hour timecourse (% condensed nuclei 70 min post-OptoBax activation: Jasplakinolide, 12% condensed nuclei (14/115 total cells); Latrunculin A, 5% condensed nuclei (8/148 total cells); No treatment control, 55% condensed nuclei (54/98 total cells); Supporting Figure 2). Finally, the actin to mitochondria translocation event occurred as previously described in the no treatment cells, yet was infrequently observed with the jasplakinolide (3%; 3/115 total cells) and latrunculin A (1%; 2/148 total cells)-treated populations post-OptoBax activation. We note that while these compounds delayed apoptotic onset in our system, both compounds have been demonstrated to be pro-apoptotic in cell culture over lengthy incubation periods, precluding lengthier experiments[34–36]. Nonetheless, it is clear that perturbations of actin dynamics delay key components of our light-activated apoptotic cascade.

It is also important to note that our observed correlation between Bax-induced MOMP and actin rearrangement may not be entirely causal. For instance, it may be that the rapid mitochondrial depolarization induced by small molecule-initiated apoptosis (i.e. staurosporine (STS)) triggers events that lead to actin redistribution in advance of endogenous BAX translocation. For example, in STS-induced apoptosis, mitochondrial depolarization occurs rapidly after treatment, while BAX maximally translocates to the OMM several hours post-STS treatment[37, 38]. As a result, in our system, we cannot rule out the possibility that BAX dependent mitochondrial depolarization drives actin redistribution via a partially MOMP-independent mechanism. In addition to actin redistribution, we also observed the nuclear accumulation of GFP-Actin in the wake of OptoBAX activation, occurring just after the appearance of mitochondrial GFP-Actin (Fig. 3C). The transition of cytosolic actin to nuclear actin has been previously described, and may be part of a general cellular stress response.[39, 40] The question of whether this freshly-nuclear localized actin binds to chromatin may also have interesting implications for actin’s role in the repression or activation of transcription of genes vital to completing the apoptotic cascade in mammalian cells, reinforcing the notion of actin as an important part of the apoptotic transcriptome, as previously observed in yeast [41–44]. Furthermore, redistribution of cytoskeletal actin into the nucleus may also provide insight into the timing of disruption of the nuclear-cytoplasmic barrier – a key apoptotic event that precipitates many of the later events in apoptosis by providing access of cytoplasmic proteins to the nuclear compartment[45–47]. Finally, increased chromatin condensation (Fig. 3D) was observed concurrently with an increase in phosphatidylserine inversion (Fig. 3E), both events peaking 70 – 90 minutes post OptoBax activation. In our next multi-color experiment, we utilized a marker of caspase cleavage to observe the downstream cleavage of caspase 3/7 (Fig. 4A). We observed initial caspase cleavage beginning within 1 hour of BAX recruitment, and increasing steadily thereafter (Fig. 4B). As anticipated, this event begins early, but peaks much later (> 3h post-BAX recruitment), consistent with previous western blotting assessment of caspase cleavage in the OptoBAX system.[16] By contrast, control experiments utilizing Cry2(1-531).L348F.mCh and Tom20.CIB did not activate the caspase activity indicator dye or the marker for phosphatidylserine inversion within the same time period of recruitment to the OMM and imaging (Supporting Figure 3). Interestingly, while we observe chromatin condensation during the course of our experiments, we did not observe significant nuclear fragmentation during the same timeframe (<1% of cells over a 120 min timecourse), suggesting that, like peak caspase cleavage, the nuclear fragmentation event also occurs later in Bax-mediated apoptosis. As a corollary to these results, longer timecourse experiments in HeLa cells utilizing a programmable LED device for cellular illumination, followed by nuclear staining and immunostaining, demonstrated nuclear fragmentation approximately 7 hours post-apoptotic induction (Supporting Figures 4 and 5). Utilizing the data from these experiments we assembled a timeline of these early apoptotic events relative to OptoBax activation (Fig. 5). This timeline demonstrates that much happens within the first two hours of the apoptotic cascade, with certain events appearing to be synchronized. For example, the actin cytoskeleton, long implicated as a key player in apoptotic progression, undergoes redistribution from the cytosol to the mitochondria in addition to movement from the cytosol to the nucleus.

This redistribution event generally precedes phosphatidylserine inversion. It is possible that wholesale redistribution of actin, a key structural component of the cytoskeleton, contributes to membrane instability and subsequent inversion. Interestingly, the increase in accumulation of Annexin V is also concurrent with an increase in chromatin condensation. These events (chromatin condensation and Annexin V accumulation) occur within the same timeframe as actin redistribution, implicating collapse and rearrangement of the actin cytoskeleton as a common thread between these events in the progression of apoptosis[28]. Finally, previous studies of staurosporine-induced apoptosis have reported that actin redistribution to the OMM occurs well before BAX translocation. In our system, recruitment of OptoBax to the OMM precipitates the actin-OMM localization event. Staurosporine has been shown to effect multiple apoptotic pathways (intrinsic and extrinsic), which may be the basis for the timing of the BAX localization event relative to that of actin.[48] Utilization of an optogenetic system such as OptoBax for the control of a single event in the apoptotic cascade may provide a more precise way for examining these translocation events. For example, other studies have focused on defining the pro-apoptotic role of cofilin at the OMM[49], yet the universal importance of the role played by cofilin and other ABPs at the OMM during apoptosis remains a matter of debate[29]. In future studies, we will attempt to resolve the complicated web of translocation events at the OMM occurring in early apoptosis using our recently developed optogenetic tools.

**Figure 3.**
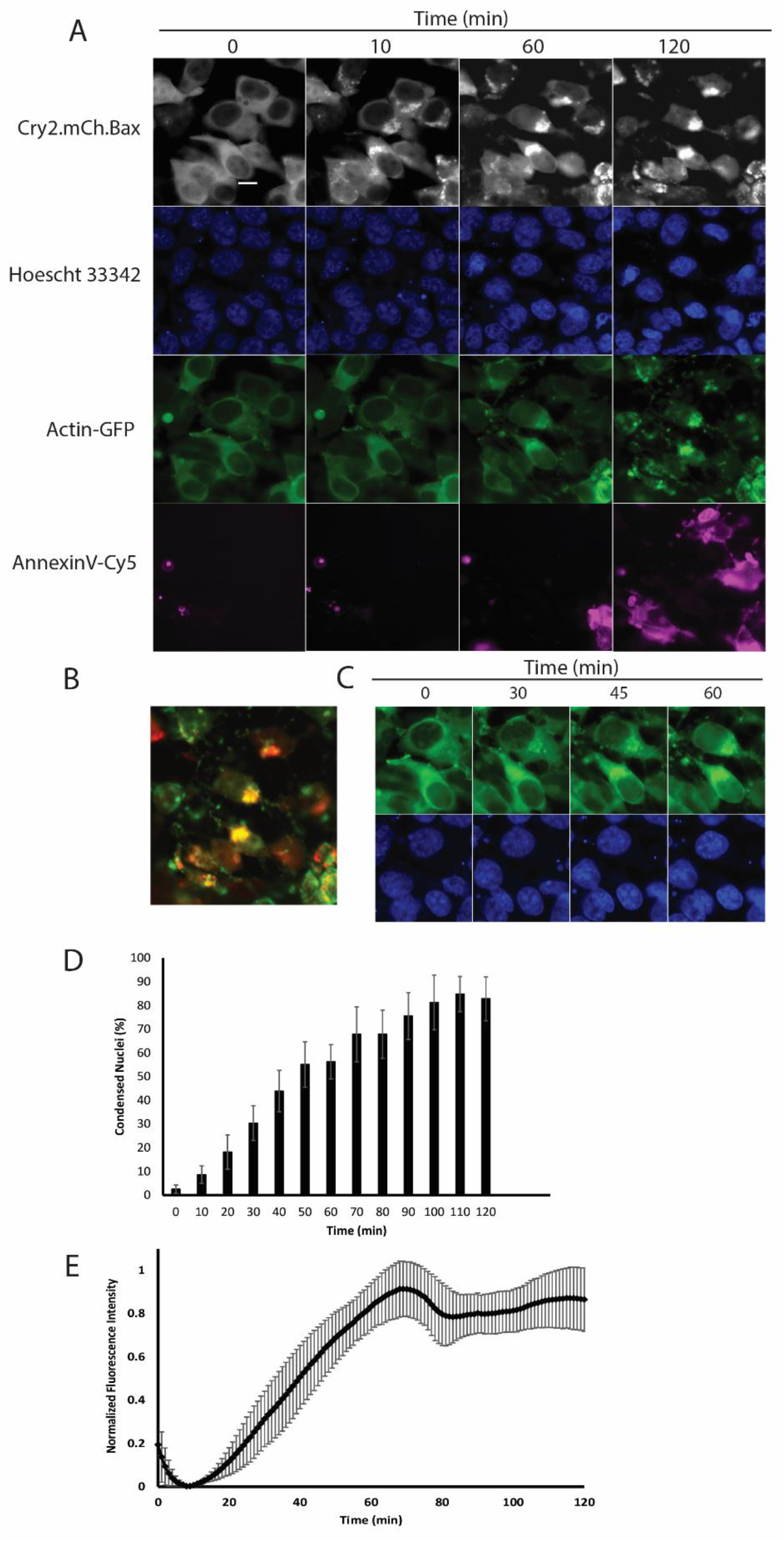
FOUR COLOR EXPERIMENT. A. Multi-color imaging (Leica DmI8 widefield microscope equipped with OKOLab stage-top incubator) of mitochondrial recruitment, chromatin condensation, actin rearrangement, and phosphotidylserine exposure in HEK293T cells. B. Overlay of actin/mitochondria after 2 hours of imaging. C. Timelapse of actin rearrangement and accumulation at mitochondria. D. % chromatin condensation versus time for chromatin marker (Hoescht 33342) and E. fluorescent intensity changes for PI marker (Annexin V-Cy5). Images acquired every minute over a two hour time course. Scale bar = 10 microns.

**Figure 4.**
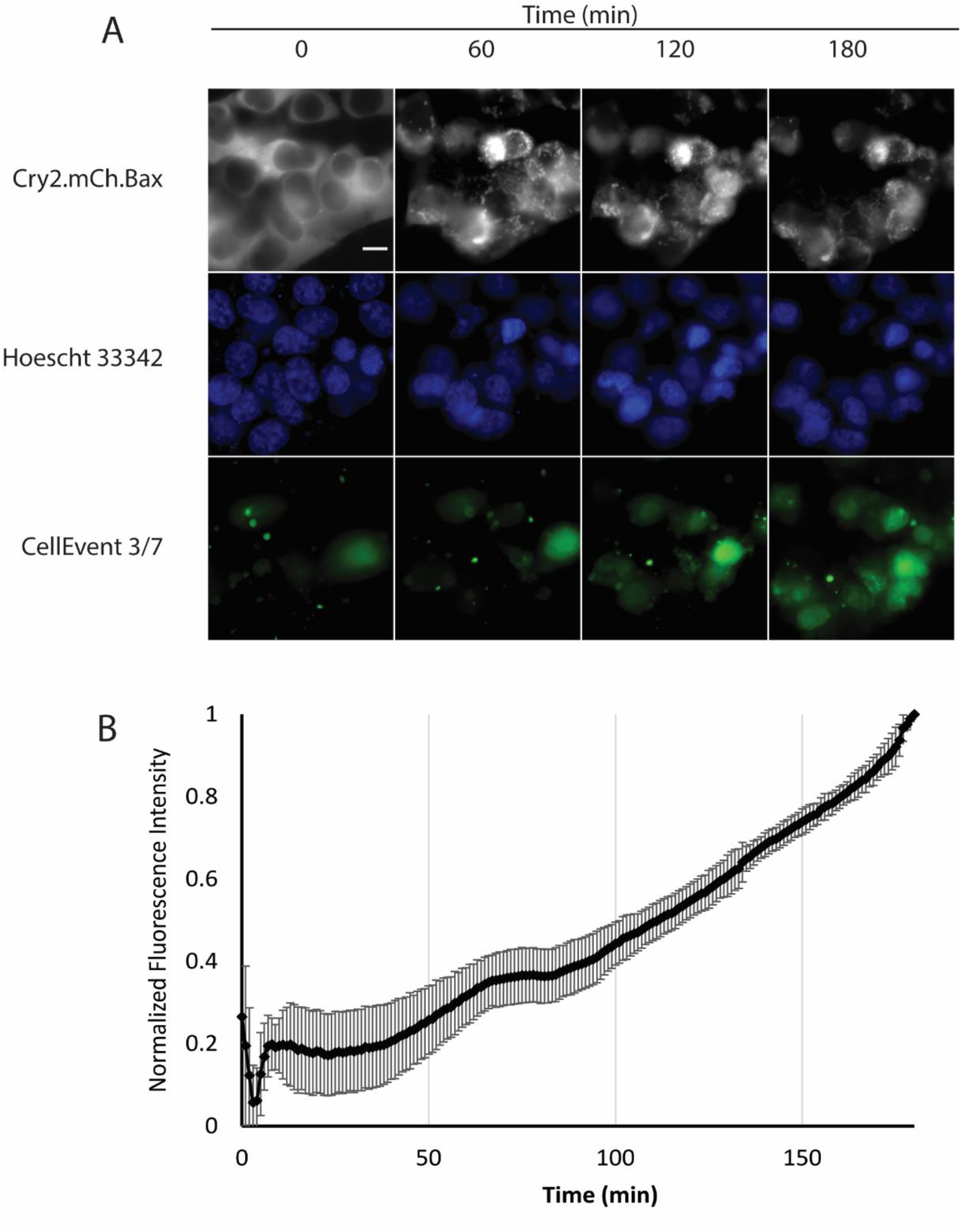
THREE COLOR EXPERIMENT. A. Multi-color imaging (Leica DmI8 widefield microscope equipped with OKOLab stage-top incubator) of mitochondrial recruitment, chromatin condensation, caspase cleavage in HEK293T cells. B. Fluorescence intensity versus time for caspase activity marker (CellEvent 3/7). Images acquired every minute over a two hour time course. Scale bar = 10 microns.

## Materials and Methods

### Cloning

The Cry2(1-531) gene was amplified by polymerase chain reaction (NEB Q5 polymerase) from the full length Cryptochrome-2 gene using a forward primer encoding an XhoI-restriction site with the sequence: GGCCAACTCGAGATGAAGATGGACAAAAAGAC and a reverse primer encoding an XmaI restriction site with the sequence: TGATATCCCCGGGCTACTTGTTGGTCATTAGAAG. The resulting PCR amplicon was then digested, gel purified, and ligated into a mCherry-N1 vector (Clontech) containing BAX S184E as a fusion to the C-terminus of mCherry. Site directed mutagenesis was used to introduce the L348F mutation using a forward primer with sequence: CCGGAATGAGAGAGTTTTGGGCTACCGGATGG; and reverse primer with the sequence: CCATCCGGTAGCCCAAAACTCTCTCATTCCGG. Cloning methods for constructs used in prior work (Cry2PHR.mCh.BAX, Tom20.CIB.GFP, and Tom20.CIB.STOP) have been described elsewhere.[16] The actin-GFP (pCAG-mGFP-actin; Addgene #21948) construct was a generous gift from Ryohei Yasuda[27].

### Cell culture and transfection

Midi prep quantities of DNA of each construct were then created from *E. coli* and collected for cell transfection. Transfections were then performed with the Lipofectamine 3000 reagent (Invitrogen) following manufacturer’s suggested protocols. Briefly, for dual transfections in 35 mm glass bottom dishes, plasmid DNA was combined in a 1:1 ratio (1,250 ng per plasmid) in 125 uL of Opti-Mem, followed by the addition of 5 uL of P3000 reagent. In a separate vial, 3.75 uL of Lipofectamine 3000 were added to 125 uL of Opti-Mem. The two 125 uL solutions were combined and allowed to incubate at room temperature for 10 min, followed by dropwise addition to cell culture. For triple-transfections, plasmid DNA was combined in a 1:1:1 ratio (1,000 ng per plasmid), followed by an identical transfection procedure. Transfection solutions were allowed to remain on cells overnight. Cells were maintained at 37°C and 5% CO2 in a humidified tissue culture incubator. HeLa and HEK293T cell lines were maintained in DMEM supplemented with 10% FBS and 1% Pen-Strep. Neuro-2a cells were maintained in Eagle’s MEM supplemented with 10% FBS and 1% Pen-Strep.

### Cell viability measurements

Background cell death experiments were performed to determine the cell toxicity of OptoBAX constructs in the absence of light exposure. Transfections were performed on HeLa cells in 6-well plates, then maintained in complete darkness for two days prior to incubation with a cell viability stain (Live-or-Dye 488/515 (Biotium). Briefly, cells were washed 1X with DPBS containing calcium and magnesium, followed by incubation at room temperature with the viability stain (diluted 1:1000 in DPBS) for 30 min. Afterwards, cells were fixed with 4% PFA in PBS for 15 min at room temperature, then washed 3 × 1 mL with DPBS, and maintained in DPBS for subsequent imaging. Images of fixed cells were acquired on a Leica Dmi8 fluorescent microscope.

### Application of fluorescent dyes

Four-color experiment: Triple-transfected cells (Cry2.mCh.Bax/Tom20.Cib /GFP-Actin) in a 35 mm glass bottom dish (Mattek) were removed from incubator, cell culture media removed by pipette, and cells gently washed 1X with DPBS supplemented with calcium and magnesium. 1 mL of PBS containing Hoescht 33342 was then applied to the cells, which were returned to the incubator for 10 min. After the elapsed time, media was removed and cells were washed with DPBS (2 × 1 mL). A pre-warmed buffer containing DPBS supplemented with 10% FBS and Annexin V-Cy5 (10 μL/1 mL of buffer; ThermoFisher A23204) were added to cells, which were promptly placed on the microscope for initiation of the light-activated experiment. Three-color experiment: Dual-transfected cells (Cry2.mCh.Bax/Tom20.Cib.STOP) were removed from incubator, cell culture media removed by pipette, and cells gently washed 1X with DPBS supplemented with calcium and magnesium. 1 mL of PBS containing Hoescht 33342 was then applied to the cells, which were returned to the incubator for 10 min. After the elapsed time, media was removed and cells were washed with DPBS (2 × 1 mL). A pre-warmed buffer containing DPBS supplemented with 10% FBS and CellEvent Caspase 3/7 Green (1 μL/1 mL of buffer; ThermoFisher C10427) were added to the cells, which were promptly placed on the microscope for initiation of the light-activated experiment.

### Cell treatments with actin binding compounds

Cells were pre-treated with actin-binding compounds for one hour prior to imaging. A. Jasplakinolide (Santa Cruz Biotechnology) was diluted from a 1 mM stock in DMSO to a working concentration of 200 nM in DPBS supplemented with 10% FBS. Prior to imaging, the cells were incubated with Hoescht 33342 for 10 minutes, followed by a 1 mL PBS wash. Cells were then resupplied with media for live cell imaging (PBS/10%FBS/200 nM Jasplakinolide/Annexin V-Cy5). B. Latrunculin A (Santa Cruz Biotechnology) was diluted from a 1 mM stock in DMSO to a working concentration of 2 μM in DPBS supplemented with 10% FBS. Prior to imaging, the cells were incubated with Hoescht 33342 for 10 minutes, followed by a 1 mL PBS wash. Cells were then resupplied with media for live cell imaging (PBS/10%FBS/2 μM Latrunculin A/Annexin V-Cy5).

### Microscopy

Leica DMi8 Live Cell Imaging System, equipped with an OKOLab stage-top live cell incubation system, LASX software, Leica HCX PL APO 63×/1.40-0.60na oil objective, Lumencor LED light engine, CTRadvanced+ power supply, and a Leica DFC900 GT camera, was used to acquire images. Exposure times were set at 150 ms (GFP, 470 nm), 150 ms (DAPI, 395 nm), 300 ms (mCherry, 550 nm), and 300 ms (Cy5, 640 nm), with LED light sources at 50% power, and images acquired every minute or every 10 minutes.

### Microscopy data analysis

Analysis of microscopy date was performed in FIJI equipped with the BioFormats package. Fluorescence intensity measurements were normalized on a scale from 0 – 1 prior to being averaged. Average normalized intensities were computed from 4 – 6 fields of view on a 63× objective, with each field of view encompassing 100 (+/-10) transfected cells. Mitochondrial residence time plots were generated by quantifying the change in fluorescence intensity at the mitochondria over time, post light recruitment. Fluoresence intensity measurments were normalized and averaged from 6 mitochondrial clusters per transfection condition. After normalizing the change in fluorescence over time, the mitochondrial association half-life was defined as the time for fluorescence intensity of the light recruited reagent to reach 50% maximum intensity at the mitochondria. Image overlays were created using the Merge Channels function in FIJI.

### Western blot

HeLa cells were lysed post-transfection with 200 µL of M-PER lysis buffer (Thermo Scientific) plus protease inhibitors. After 10 min on a rotary shaker at room temperature, lysates were collected and centrifuged for 15 min (1000 rpm at 4 °C) and supernatants reserved. The resulting lysates were subjected to electrophoresis on a 10% SDS-PAGE gel and then transferred to PVDF membranes (20 V, overnight, at 4 °C). Membranes were then blocked for 1 h with 5% BSA in TBS with 1% Tween (TBST), followed by incubation with the primary antibody (Rockland Anti-mCherry antibody 600-401-P16) 1:1000 dilution in 5% milk – TBST overnight at 4 °C on a platform rocker. The membranes were then washed 3 × 5 min each with TBST and incubated with anti-rabbit IgG – HRP secondary antibody (abcam 6721) at 1:3000 in 5% BSA – TBST for 2 h at room temp. After washing 3 × 5 min with TBST, the membranes were exposed to a chemiluminescent substrate for 5 min and imaged with using an Azure cSeries imaging station.

### Programmable LED device

The device (designed and constructed by Hoffman and Hughes) combines four wavelength ranges (440-460nm, 520-540nm, 650-670nm, 720-750nm) into a single package using Surface Mount Technology (SMT) and is powered with an Arduino UNO system. Surface Mount Devices (SMD) Luxeon Rebel and Luxeon Color series LEDs were soldered on 20mm mounts with a hot air gun designed for Surface Mount Devices. The LEDs/MCPCB aluminum LED bases were affixed to a 130-mm × 70 mm Rectangular 15 mm high heat sink to prevent overheating and allow for longer exposures of samples. The intensity of the light was controlled with a combination of variable resistors, PNP transistors, and zener diodes. The LED arrays are controlled using an Arduino controller to control illumination, time, intensity, and wavelength.

### Cellular illumination and immunostaining

HeLa cells were plated in 35 mm glass bottom dishes (Mattek) at a density of 2.0 × 10^5^ cells per dish and maintained at 37 °C in a humidity-controlled incubator with a 5% CO_2_ atmosphere in DMEM (10% FBS, 1% Pen-Strep). The following day, cells were transfected with Cry2.L348F.531.mCh.BAX and Tom20.CIB.GFP as previously reported. 24 h post transfection, cells were illuminated with 440-460 nm light using a programmable LED device prior to being returned to the incubator. At the conclusion of the dark incubation period, cells were washed 3 × 1 mL with PBS and then fixed with 1 mL of 4% paraformaldehyde in PBS at room temperature for 10 min. Cells were washed 2 × 1 mL with PBS and blocked and permeablized for 20 min in antibody dilution buffer (1% BSA; 0.3% Triton-X-100; PBS) at 4 ºC. Blocking was followed by overnight incubation at 4 ºC with rabbit anti-cleaved caspase 3 antibody (Cell Signaling 9661) at 1:400 dilution in antibody dilution buffer (1% BSA; 0.3% Triton-X-100; PBS). Cells were then washed with PBS (3 × 5 min) before incubation with anti-rabbit AlexaFluor 647 secondary antibody (Life Technologies A21245) at 1:500 dilution in antibody dilution buffer. After washing cells with PBS (3 × 5 min), nuclei were stained briefly (5 min) with Hoescht 33342 (1 μg/mL), followed by image acquisition on a confocal microscope (Olympus IX2-DSU Tandem Spinning Disk Confocal; 60X objective).

### Plasmids

All plasmids generate for this publication will be available through Addgene, Inc., a not-for-profit plasmid DNA repository.

## Acknowledgements

The authors wish to thank Dr. Karen Litwa (ECU Brody School of Medicine) for the gift of HEK293T cells; Dr. Alex Murashov (ECU Brody School of Medicine) for the gift of Neuro-2a cells; Dr. Nathan Hudson (ECU Department of Physics) for use of the Leica Dmi8 widefield microscope.

---

**SCHEME 1:**
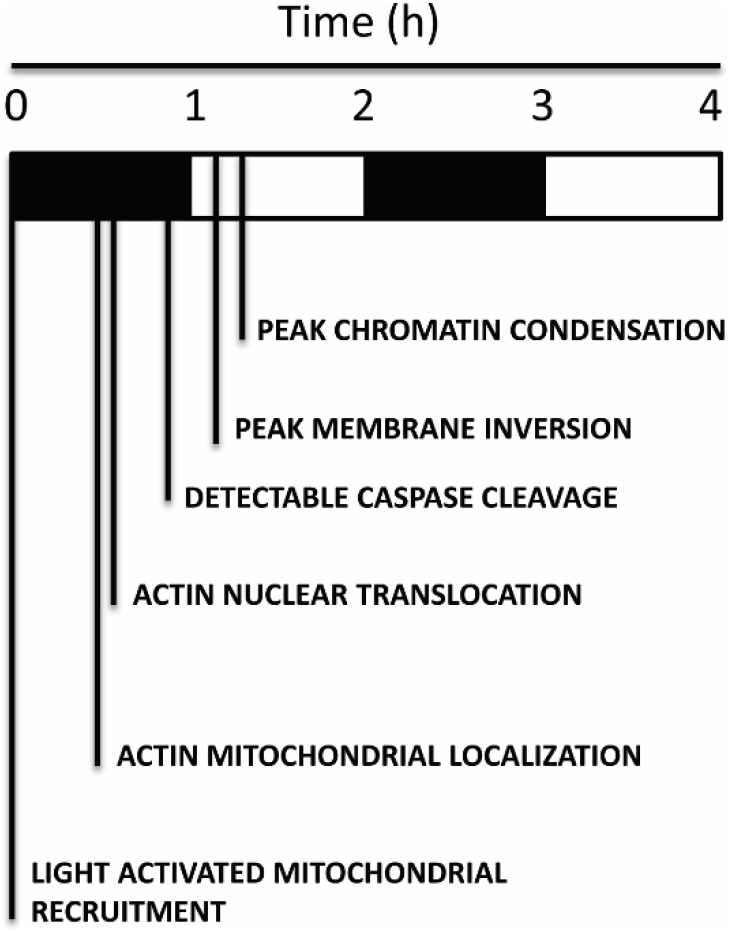
TIMELINE OF APOPTOTIC EVENTS.

**Supporting Figure 1.**
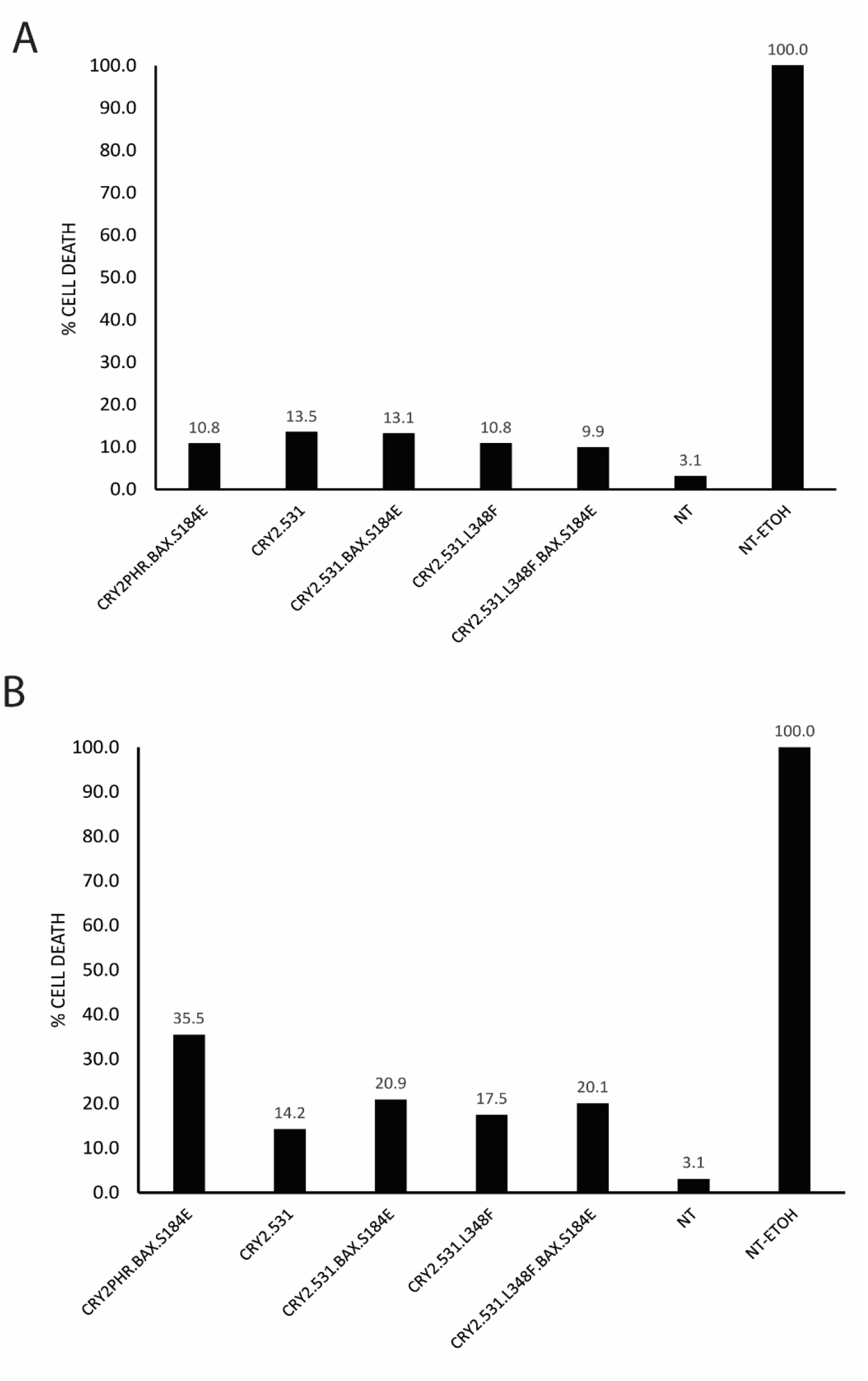
ASSESSMENT OF BACKGROUND TOXICITY. A. Quantification of Biotium Live-Or-Dye staining of singly transfected HeLa cells. NT = no transfect; NT-ETOH = no transfect, 10 min treatment with 15% EtOH. >500 cells were counted for each experimental condition on a widefield fluorescent microscope. B. Quantification of Biotium Live-Or-Dye staining of double transfect HeLa cells (Cry2 variant + Tom20.Cib). NT = no transfect; NT-ETOH = no transfect, 10 min treatment with 15% EtOH. >500 cells were counted for each experimental condition on a widefield fluorescent microscope.

**Supporting Figure 2.**
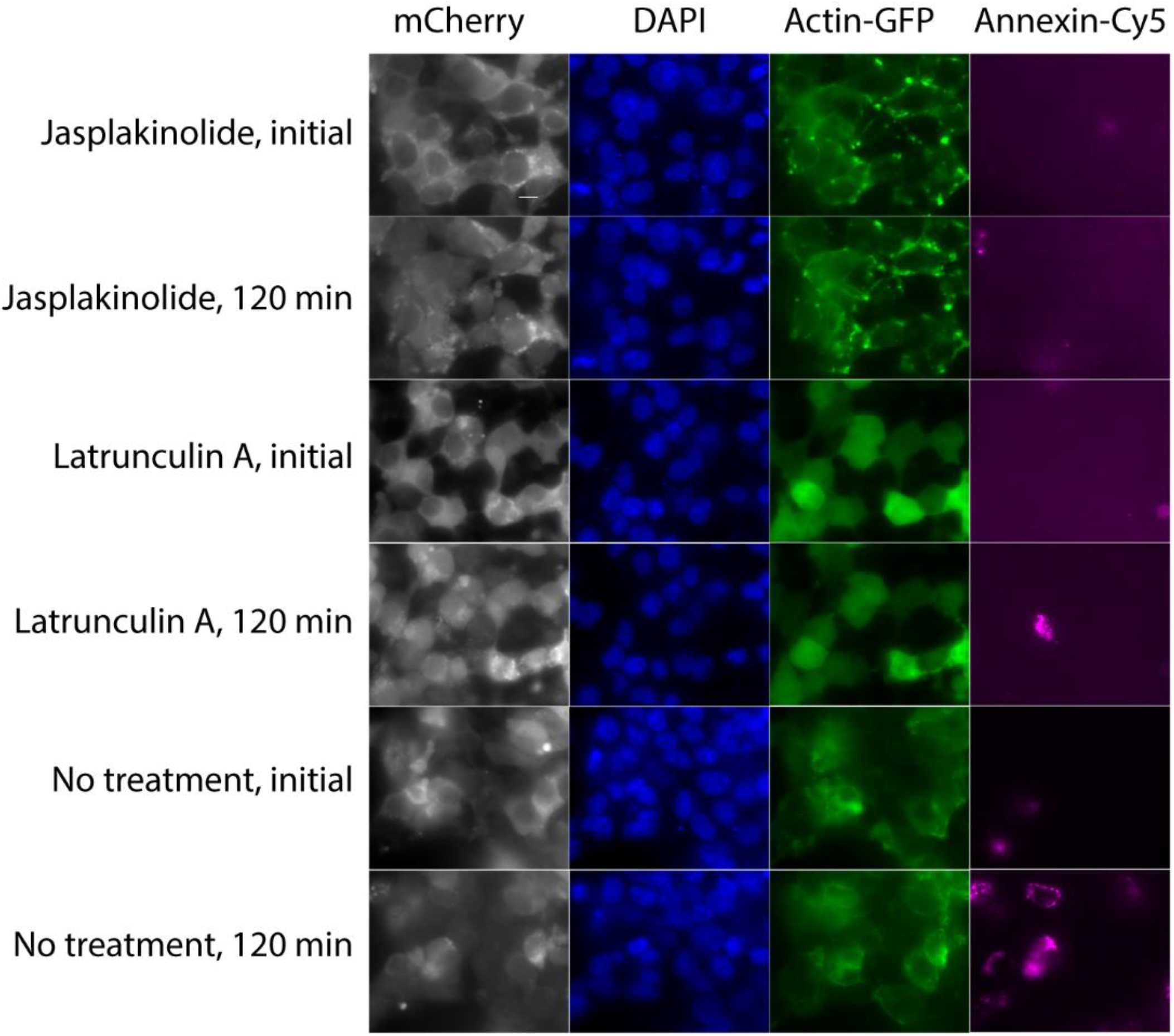
Pharmacological Inhibition of Actin Dynamics. Time points from multi-color imaging of HEK293T cells transfected with Cry2.531.L348F.mCh.BAX.S184E /Tom20.Cib and incubated with cell permeable inhibitors of actin dynamics (200 nM Jasplakinolide; 2 μM Latrunculin A). Images acquired every minute over a two hour time course. Scale bar = 10 microns.

**Supporting Figure 3.**
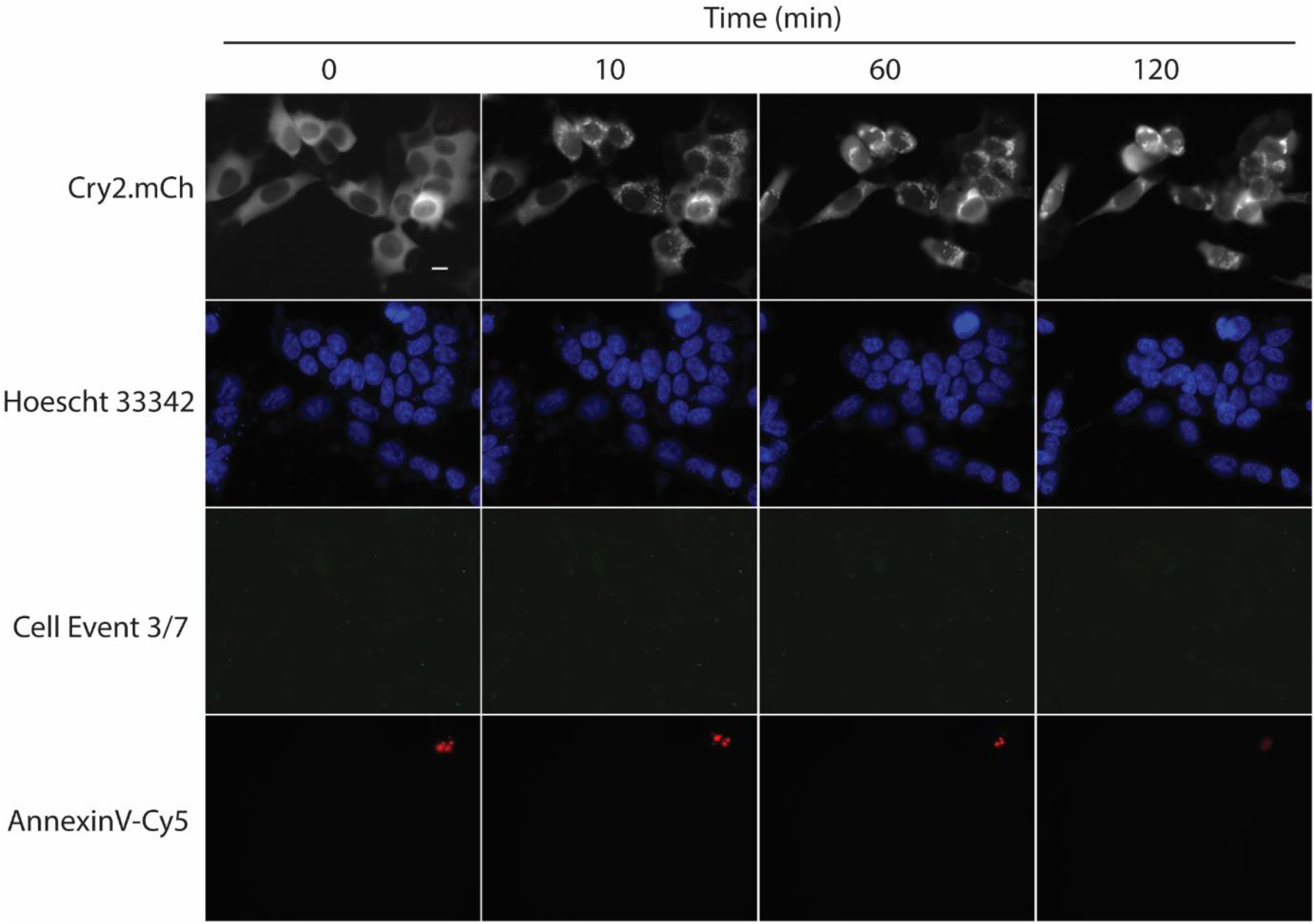
Cry2(1-531).L348F.mCh/Tom20.Cib Control Experiment. Time points from multi-color imaging of HEK293T cells transfected with Cry2.531.L348F.mCh/Tom20.Cib and incubated in the presence of a marker for phosphatidylserine inversion (Annexin-Cy5) and caspase cleavage (CellEvent 3/7 Green). Images acquired every minute over a two hour time course. Scale bar = 10 microns.

**Supporting Figure 4.**
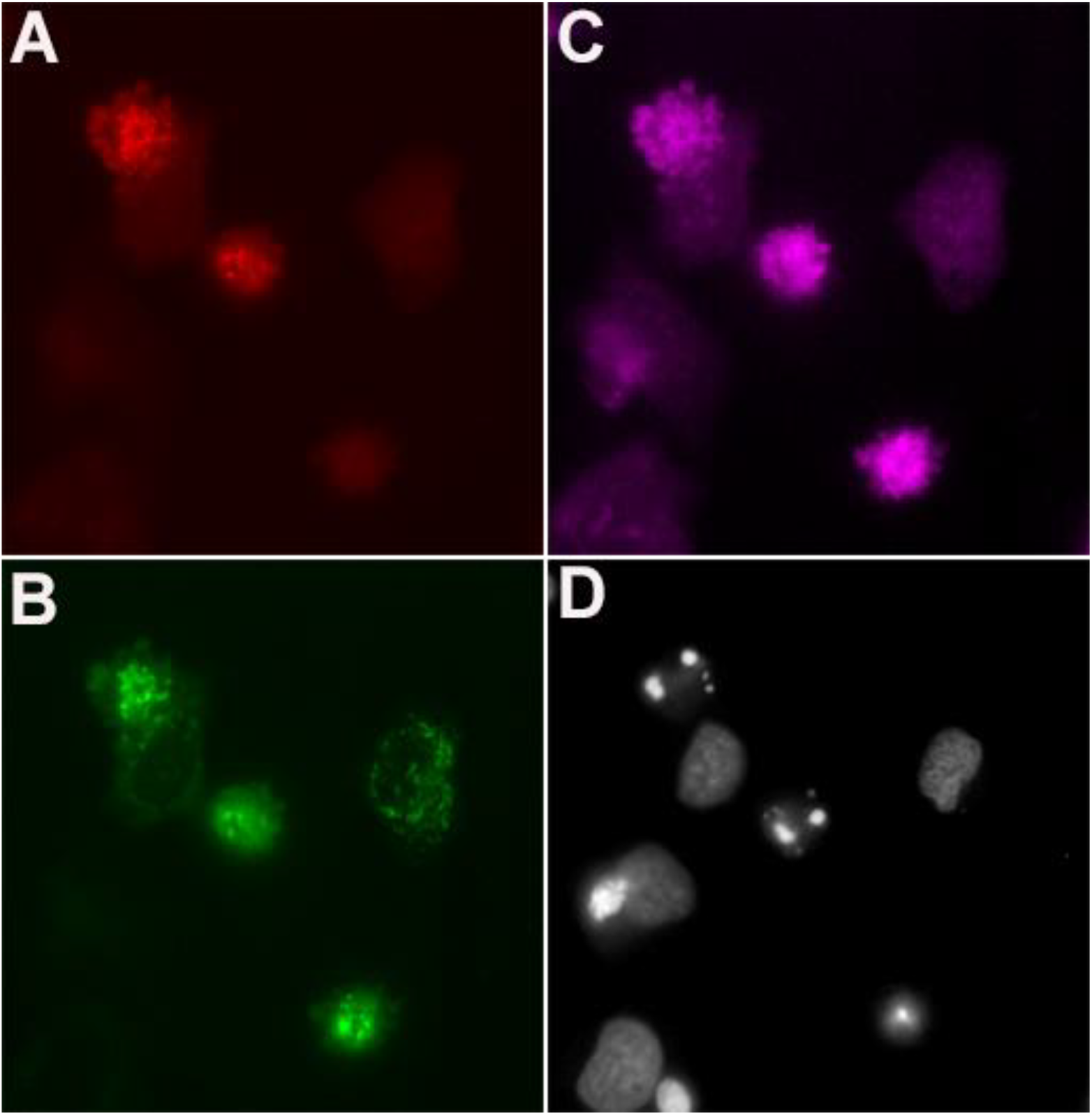
Immunostaining of light-treated HeLa cells 7 h post-illumination. HeLa cells exposed to 440-460 nm blue light (1 pulse/sec; 5 min total light exposure), followed by fixation and immunostaining. A. Cry2.L348F.531.mCh.BAX (mCh fluorescence) B. Tom20.CIB.GFP (GFP fluorescence) C. α-cleaved caspase 3/7 fragment (Cy5 fluorescence) D. Hoescht 33342 stained nuclei (DAPI fluorescence).

**Supporting Figure 5.**
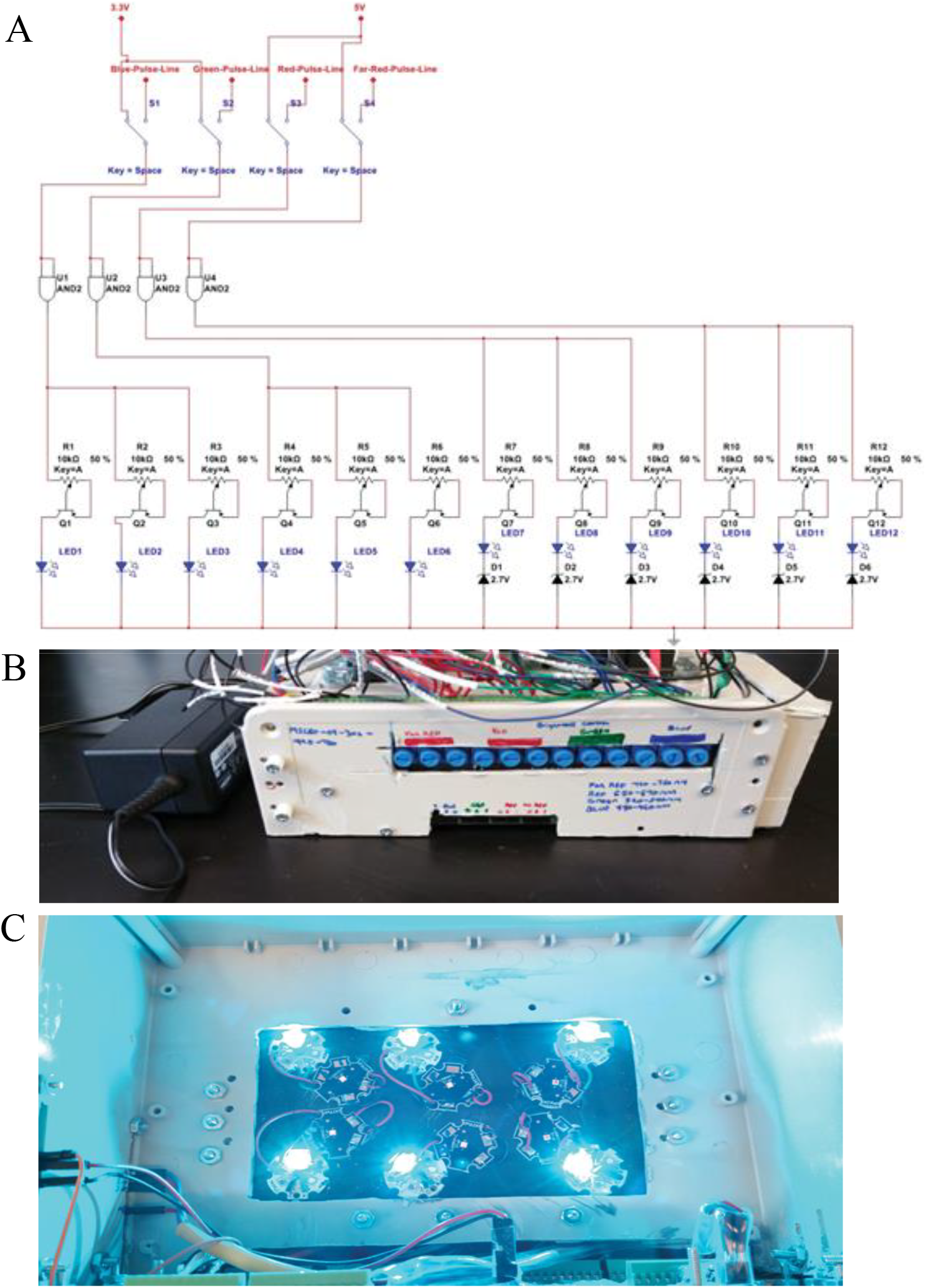
Schematic of Programmable Multi-Wavelength LED Device. A. Circuit diagram of programmable LED device. B. Side view of light intensity control module of programmable device. C. 440-460 nm LEDs illuminated at full power.

**Supporting Movie 1.** Movie of Cry2(1-531).mCh vs. Cry2(1-531).L348F.mCherry recruitment to the OMM (Tom20.CIB.GFP) in HeLa cells. Images were acquired every 30 s; recruitment to OMM initiated with a single 150 ms pulse of 470 nm light.

**Supporting Movie 2.** Movie of Cry2(1-531).L348F.mCh.BAX.S184E vs. Cry2(1-531).L348F.mCherry recruitment to the OMM (Tom20.CIB.GFP) in HeLa cells. Images were acquired every 10 min over a 2 h time course; recruitment to OMM sustained with one 150 ms pulse of 470 nm light every 10 min.

**Supporting Movie 3.** Multi-color imaging of apoptotic markers during a 2 h timecourse of recruitment and imaging in HEK293T cells. Panels depict (clockwise, starting from upper left): Cry2(1-531).L348F.mCh.BAX.S184E (mCherry fluorescence); Hoescht 33342 (DAPI fluorescence); Annexin V-Cy5 (Cy5 fluorescence); GFP-Actin (GFP fluorescence). Images acquired every 1 min.

